# Urbanisation drives non-linear, guild-specific restructuring of fungal communities in coastal dune habitats

**DOI:** 10.64898/2025.12.11.693695

**Authors:** Laertis Bechlis, Thom Stoppelenburg, Jan-Berend Stuut, Willem Renema, S. Emilia Hannula, Jan-Niklas Macher

## Abstract

Coastal dunes form dynamic ecosystems where fungal communities play critical roles in nutrient cycling and plant establishment. Urban expansion increasingly fragments these habitats through trampling, beach grooming, and nutrient inputs, yet the effects of urbanization on below□ground fungal diversity remain poorly understood. We used eDNA metabarcoding on 294 soil cores from 49 transects spanning 35 km of the Dutch coast to test how urban proximity influences total fungal diversity and composition in different functional guilds (litter saprotrophs, plant pathogens, mycorrhizal fungi) across incipient (early-successional) and established dunes. Urban effects were habitat-specific and generally strongest in incipient dunes. In incipient dunes, total alpha diversity showed a U-shaped relationship with city distance, with minima at intermediate distances, whereas established dunes showed weak responses. Fungal guilds diverged in their responses as predicted: litter saprotroph diversity increased with distance from cities in both habitats (consistent with greater organic inputs by vegetation), plant pathogens showed U-shaped diversity in incipient dunes, suggesting distinct urban-associated and native-associated assemblages, and mycorrhizal diversity increased with distance in both habitats, consistent with limited suitable hosts and higher disturbance near cities. Partitioning of beta diversity revealed that turnover, rather than nestedness, accounted for almost all variation in community composition, indicating taxon replacement rather than directional loss. Community uniqueness was higher near cities, particularly in incipient dunes and for most guilds, implying that urban environments foster compositionally atypical assemblages relative to the regional pool. Overall, urbanization acts as a trait-based environmental filter, restructuring fungal communities without necessarily reducing fungal diversity. Stronger responses in early-successional dunes highlight their vulnerability to combined natural and urban stressors and the need to prioritize protection of incipient coastal habitats under intensifying urban pressure.

## Introduction

Coastal sand dunes form dynamic ecosystems shaped by geomorphological processes, harsh environmental conditions and ecological succession (Hesp, 1991; Anwar Maun, 2009; McLachlan and Defeo, 2017). Species inhabiting these ecosystems at the terrestrial-marine boundary experience shifting sand, periodic flooding, salt spray, and low nutrient availability, which creates strong abiotic gradients from the sea to inland (Miller et al., 2010; Roy-Bolduc et al., 2016a). Along these gradients, vegetation transitions from sparse, stress-tolerant pioneer species in incipient dune habitats to more diverse and structurally complex plant assemblages in stabilised, established dunes. Urban expansion increasingly encroaches on dunes worldwide, fragmenting habitats via trampling, beach grooming and shoreline engineering that remove wrack and organic inputs, and via nutrient and pollutant inputs, microclimatic change, and plant invasions, all of which have strong impacts on biodiversity (Kutiel et al., 2000; Parsons et al., 2020). These urban stressors alter soil microbial communities and ecosystem functioning (Cardinale et al., 2012; Novoa et al., 2020), can reshape biodiversity, and impact ecosystem services such as shoreline protection (Carranza et al., 2020; Aguilera et al., 2022).

Although plant and animal responses to urbanisation in dune ecosystems have been studied (Zarnetske et al., 2010; Bouziane et al., 2020; McGuirk et al., 2022), belowground microbes remain understudied despite their key roles in nutrient cycling, decomposition, soil structure, and plant dynamics (van der Heijden et al., 1998; Bardgett and van der Putten, 2014; Tedersoo et al., 2014; Bahram and Netherway, 2022). In dunes, fungi mediate the decomposition of scarce organic inputs, enhance soil fertility under oligotrophic conditions, and facilitate plant establishment through mutualisms such as mycorrhizal symbioses (Koske and Gemma, 1997; Prenafeta Boldú et al., 2014; da Silva et al., 2015; Roy-Bolduc et al., 2016b). Fungal communities are therefore linked to vegetation, and plant richness and diversity can influence fungal richness and guild composition through host availability and resource inputs. Across the succession gradient from incipient to established dunes, fungal communities and guilds shift with incipient dunes hosting more stress-tolerant, generalist saprotrophs, whereas established dunes support richer assemblages including specialised litter decomposers and plant-associated taxa (Prenafeta Boldú et al., 2014; Roy-Bolduc et al., 2015; Cao et al., 2024) Yet recent work shows that incipient dunes can also harbor unexpectedly rich microbial assemblages, including marine and terrestrial taxa that can handle the stress of this environment (Nayak et al., 2019; Spaeth et al., 2024).

Despite this background, we still lack a clear picture of how urbanisation modifies these successional patterns. Urban drivers in dunes operate through multiple pathways, such as reduced organic matter inputs due to beach cleaning (limiting saprotrophic resources), physical disturbance from recreation and infrastructure (disrupting hyphal networks), nutrient enrichment and pollutants (shifting competitive hierarchies), microclimatic change, host-plant turnover or plant richness via landscaping and the occurrence of ornamental plants (Parsons et al., 2020; Liu et al., 2023; Khalid et al., 2024). Because these pressures act on top of strong natural stressors in these dynamic systems, responses are expected to depend on habitat context and fungal guild traits: early-successional, resource-limited incipient dunes should be more sensitive to urbanisation effects, whereas established dunes with developed soils and vegetation may buffer impacts.

Different fungal guilds are expected to react in different ways to urban impacts (Hannula et al., 2017; Hannula and Träger, 2020; Rillig et al., 2023). Saprotrophs can be expected to track organic matter inputs, while patterns in mycorrhizal fungi should follow that of host plants, and plant pathogens likely track host diversity and density. Urbanisation can therefore shift fungal communities by changing resource availability, microclimate and host pools, and potentially favouring disturbance-tolerant generalists near cities and more specialised taxa in less impacted sites. Urban effects can lead to taxon replacement (turnover) or to directional loss or gain (nestedness). Disentangling these processes, and identifying where communities are atypical relative to the regional pool, is essential for interpretation of biodiversity patterns and management of ecosystems. Here we test how urbanisation affects soil fungal richness, composition, and community structure in coastal incipient and established dunes. We use soil environmental DNA (eDNA) metabarcoding with functional guild annotation (Põlme et al., 2020) to quantify responses along an urban gradient along the North Sea coastline.

We hypothesize that (1) effects of urbanisation are stronger in incipient dunes than in established dunes; (2) functional guilds (saprotrophs, mycorrhizal fungi, plant pathogens) differ in their responses to urban proximity according to their traits and ecological roles, and (3) sites near cities will show more unique community composition, consistent with urban influences such as altered host flora (e.g., ornamental plants). By resolving fungal responses across early and later dune succession stages, we provide new insight into the diversity, composition, and distinctiveness of coastal dune fungal communities, and provide evidence that can guide conservation and management of these dynamic ecosystems under intensifying urban pressure.

## Material & Methods

### Sampling

Soil samples were collected between 25 July and 14 August 2023 from 49 transects spanning a 35 km stretch of coastline between the cities of Scheveningen and Zandvoort, the Netherlands. At each transect, three samples were taken from the incipient dune zone, spaced evenly from the base to the end of the incipient dunes, which was marked by the trough separating them from the established (older) dunes. Three samples were taken from the established dunes, between the base and the ridge, where vegetation shifts from grasses and shrubs to dune heath. In cases where the incipient dunes were heavily degraded due to buildings or heavy trampling within the dune area, sampling locations were chosen based on the position and extent of the nearest remaining incipient dunes. Samples were collected using sterile 20 mL plastic cores, inserted into the sediment to the 10 mL mark. The collected material was transferred into sterile 50 mL falcon tubes and stored at - 20°C until further processing. In total, 294 samples were collected.

For each transect, we measured grain size separately for the incipient dune and established dune samples. The three samples from each zone were pooled in equal volumes, and the average grain size (in μm) was determined using a LS13320 Particle Size Analyzer (Beckman-Coulter, USA). To assess distance from the next city, we measured the distance (in meters) from each transect to the nearest city center. This was defined as the distance to the main, central beach access point along the city boulevard and was measured using Quantum GIS (v3.42.1). See Supplementary Table 1 for sampling coordinates and metadata.

### DNA extraction, amplification and sequencing

DNA was extracted from soil samples using the SDE protocol (Bollmann-Giolai et al., 2020). Fungal DNA was amplified using a two-step PCR protocol, using the ITS3 forward primer mix with the ITS4ngs reverse primer (Tedersoo et al., 2014). The first PCR reaction contained 10.3 µl mQ water, 4 µl Phire Green PCR buffer (10x; Thermo fisher, Waltham, United States of America), 0.8 µl Bovine Serum Albumin (BSA, 10 mg/ml), 0.4 µl dNTPs (2.5 mM), 0.4 µl Phire II HS Taq (5U/µl), 1 µl of each nextera-tailed primer (10 pMol/µl), and 2.5 µl of DNA template. PCR amplification involved an initial denaturation at 98 °C for 30 seconds, followed by 34 cycles of denaturation for 5 seconds at 98 °C, annealing at 50 °C for 5 seconds, and extension for 15 seconds at 72 °C, concluding with a final extension at 72°C for 5 minutes. 24 negative controls (Milli-Q water, Merck, Kenilworth, USA) were processed alongside the samples, from DNA extraction to sequencing, to check for potential contamination. After the first PCR, samples were cleaned with AMPure beads (Beckman Coulter, Brea, United States) at a 0.9:1 ratio according to the protocol to remove short fragments and primer dimers. The second PCR involved amplification with individually tagged primers, following the same protocol as above and using the PCR product from the first PCR as the template, but reducing the PCR cycle number to 10. DNA concentrations were measured using the FragmentAnalyzer (Agilent Technologies, Santa Clara, CA, USA) with the DNF-910 dsDNA kit (35-1500 bp). Samples were pooled equimolarly. The final library was cleaned with AMPure beads as described above and sent for sequencing on one Illumina MiSeq run (2 × 300 bp read length) at Baseclear (Leiden, The Netherlands).

### Bioinformatic processing of community metabarcoding data

We processed the raw metabarcoding reads using APSCALE (Buchner et al., 2022) with the following settings: maximum differences in percentage: 10; minimum overlap: 50, minimum sequence length: 250 bp and the other settings, including LULU filtering, left on default. Sequences were clustered into Operational Taxonomic Units (OTUs) with a sequence similarity threshold of 97%. Following a conservative approach to ensure removal of potential low-key contamination, we conducted several steps: We subtracted the number of reads of each OTU that was found in negative controls from the read count of that OTU in each sample. Samples with less than 10 000 reads were removed from the dataset. After quality filtering, 253 samples remained in the dataset. Per sample, OTUs that did not reach a read abundance of 0.01 percent were removed. In addition, OTUs that occurred only in a single sample were removed. We performed taxonomic assignment of OTUs using the combined UNITE database (v.10, (Abarenkov et al., 2023)) containing fungi and other eukaryotes. OTUs not identified as Fungi or with <90% identity were removed. Fungi were assigned into functional guilds using the FungalTraits database (Põlme et al., 2020). Ectomycorrhizal and arbuscular mycorrhizal fungi were grouped into the category mycorrhizal. In addition, we extracted plant richness per site by counting the number of OTUs classified as Anthophyta using the UNITE database, which we used as a measure of local vascular plant richness. See Supplementary Table 2 for the full OTU table.

### Alpha Diversity

All statistical analyses were performed in R 4.4.0 (R Core Team, 2024). Analyses were run for the full fungal community and, separately, for plant pathogenic, litter saprotrophic, and mycorrhizal fungi, which we chose for their contrasting ecological roles. To assess α-diversity patterns, we computed Shannon diversity per sample after converting community data to relative abundances in the vegan package. To test effects of urbanisation and environmental variables on diversity, we fitted generalized additive mixed models (GAMMs) using mgcv, with dune type (established vs. incipient) as a categorical predictor and separate smooths of distance from the city by dune type to allow non-linear responses along the gradient. Sediment grain size and plant richness were included as additional smooth covariates. To account for the blocked sampling design, the transect was included as a random intercept. The continuous predictors (distance from city, grain size, plant richness) were z-scored. Values and 95% confidence intervals were visualised along the urbanisation gradient, and grain size and plant richness were held at their means. OTU richness was analysed with the same model structure.

### Beta Diversity

We calculated Bray-Curtis dissimilarities from OTU data after performing Hellinger transformation to stabilise variance and reduce the influence of highly abundant taxa (Legendre and Gallagher, 2001). Distance from the city centre and sediment grain size were treated as continuous variables. We performed distance-based redundancy analysis (dbRDA, in vegan) using Bray-Curtis dissimilarities as the response. To account for the blocked sampling design, we controlled for the transect by using it as a conditioning term, and permutation tests were restricted within transects (999 permutations). The explanatory variables were dune type (incipient vs. established), distance from the city, sediment grain size, the interaction dune type × distance from city, and plant richness. Because distance from a city is measured at the transect level, its main effect is not testable with the within-transect test. Results were plotted with ggplot2, with overlaying biplot arrows for predictors and 95% data ellipses for dune-type groups. To decompose multi-site β-diversity, we used betapart on presence-absence data to obtain total β (β_SOR) and its components, turnover (β_SIM) and nestedness (β_SNE).

### Local Contributions to Beta Diversity (LCBD)

We quantified per-site compositional uniqueness using Local Contributions to Beta Diversity (LCBD (Legendre and De Cáceres, 2013)). LCBD quantifies each site’s contribution to overall β-diversity, with larger values indicating more compositionally unique sites. OTU counts were Hellinger-transformed prior to analysis, and LCBD values were computed with the beta.div function in adespatial. To evaluate how LCBD varied along the urbanisation gradient, we fitted generalised additive mixed models, including dune type as a categorical variable, smooths of distance from the city within each dune type, and smooths for sediment grain size and plant richness, with transect included as a random intercept. Continuous predictors were z-scored, smoothing parameters were estimated by REML, and results with 95% confidence intervals were plotted while holding covariates of sediment grain size and plant richness at their means. Analyses were conducted for the full fungal community and separately for plant pathogenic, litter saprotrophic, and mycorrhizal fungi.

## Results

### Alpha diversity

Shannon diversity did not differ significantly between dune types (Estimate = −0.10, p = 0.30). In incipient dunes, Shannon diversity showed a clear non-linear response to distance from the city (edf = 1.96, F = 10.92, p < 0.001), whereas in established dunes diversity was unaffected by distance (edf = 1.52, F = 0.65, p = 0.59). Plant richness had a significant, slightly non-linear positive effect on Shannon diversity (edf = 1.73, F = 10.51, p < 0.001), while sediment grain size had no detectable effect (p = 0.53). The model explained 35.8% of the deviance (adjusted R² = 0.288) after accounting for transect as a random effect (Figure 1A).

**Figure 1:**
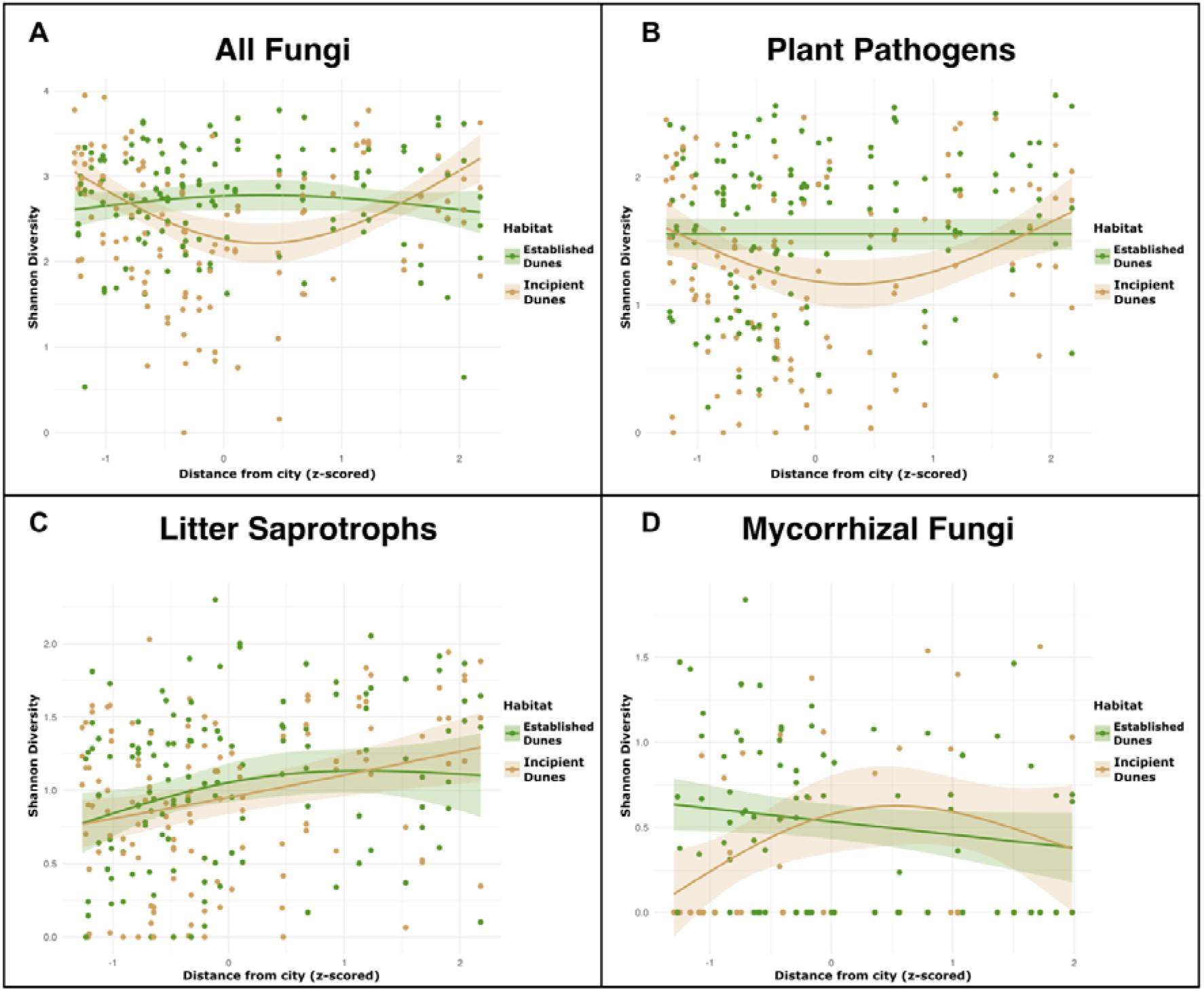
Effects of urban proximity on fungal Shannon diversity across dune habitats. Each panel shows generalized additive model (GAM) fits for Shannon diversity in relation to distance from the nearest city, with sediment grain size and plant richness held constant at their mean values. A) all fungal OTUs, B) plant pathogenic fungi, C) litter saprotrophic fungi, D) mycorrhizal fungi. Data points are shown separately for incipient dunes (ID, orange) and established dunes (ED, green), with shaded ribbons indicating 95% confidence intervals around the GAM fits.

Shannon diversity of plant pathogenic fungi was significantly higher in established dunes than in incipient dunes (Estimate for incipient vs. established dunes = −0.18, p = 0.023). In incipient dunes, Shannon diversity showed a clear non-linear response to distance from the city (edf = 1.90, F = 5.20, p = 0.0046), with diversity lowest at intermediate distances and higher both close to and far from urban areas (Figure 1B). In established dunes, Shannon diversity did not vary significantly with distance from the city (edf = 1.00, F = 1.81, p = 0.18). Plant richness had a strong, approximately linear positive effect on pathogen diversity (edf = 1.00, F = 24.71, p < 0.001), whereas sediment grain size showed no detectable effect (p = 0.48). The model explained 42.8% of the deviance (adjusted R² = 0.353) after accounting for transect as a random effect.

Shannon diversity of litter saprotrophs did not differ between established and incipient dunes (Estimate =-0.035, p = 0.625). Diversity increased with distance from the city in both dune types, with a moderate non-linear effect in established dunes (edf = 1.46, F = 4.52, p = 0.011) and a strong near-linear increase in incipient dunes (edf = 0.99, F = 7.47, p < 0.001; Figure 1C). In addition to urbanisation, sediment grain size (edf = 1.42, F = 3.84, p = 0.041) and plant richness (edf = 0.81, F = 2.55, p = 0.023) both had significant positive effects on saprotroph diversity. The model explained 24.7% of the deviance (adjusted R² = 0.184).

Shannon diversity of mycorrhizal fungi did not differ significantly between established and incipient dunes (Estimate for incipient vs. established dunes = −0.08, p = 0.43). In incipient dunes, diversity showed a moderately non-linear response to distance from the city (edf = 1.77, F = 4.99, p = 0.019), with diversity increasing along the urbanisation gradient (Figure 1D). In established dunes, mycorrhizal diversity also varied significantly with distance, with an approximately linear response (edf = 1.00, F = 4.02, p = 0.047). Neither sediment grain size (edf = 1.52, F = 1.48, p = 0.34) nor plant richness (edf = 1.00, F = 0.96, p = 0.33) had a clear effect on mycorrhizal diversity. The model explained 14.1% of the deviance (adjusted R² = 0.099) after accounting for transect as a random effect.

### Beta diversity

Distance-based redundancy analysis (dbRDA) was used to quantify how fungal community composition was shaped by dune type, urbanisation, sediment grain size, plant richness, and the interactions of grain size and dune type. To gain insight into community assembly processes, we partitioned beta diversity into turnover and nestedness components using OTU presence-absence data. Separate dbRDA models were run for the total fungal dataset and for each functional group to evaluate how these processes varied across ecological roles.

For all fungi, multi-site β-diversity partitioning (Sørensen) showed that turnover dominated total β-diversity (β_SIM = 0.986; 99.5% of β_SOR = 0.991), while nestedness contributed only 0.5% (β_SNE = 0.005). After controlling for transect, the dbRDA based on Bray–Curtis dissimilarities was significant (F = 7.99, p = 0.001). The constrained component explained 8.1% of the variation in community composition (adjusted R² = 0.089). Marginal tests indicated significant effects of plant richness (F = 4.27, p = 0.001) and of the dune type × distance-to-city interaction (F = 2.84, p = 0.001), consistent with habitat-specific responses to urbanisation, whereas sediment grain size did not explain additional variation. The first two constrained axes (CAP1 and CAP2) were highly significant (both p = 0.001) and together accounted for 94.4% of the constrained variance (75.3% and 19.1%, respectively); CAP3 was weaker but still significant (p = 0.018; Figure 2A).

**Figure 2:**
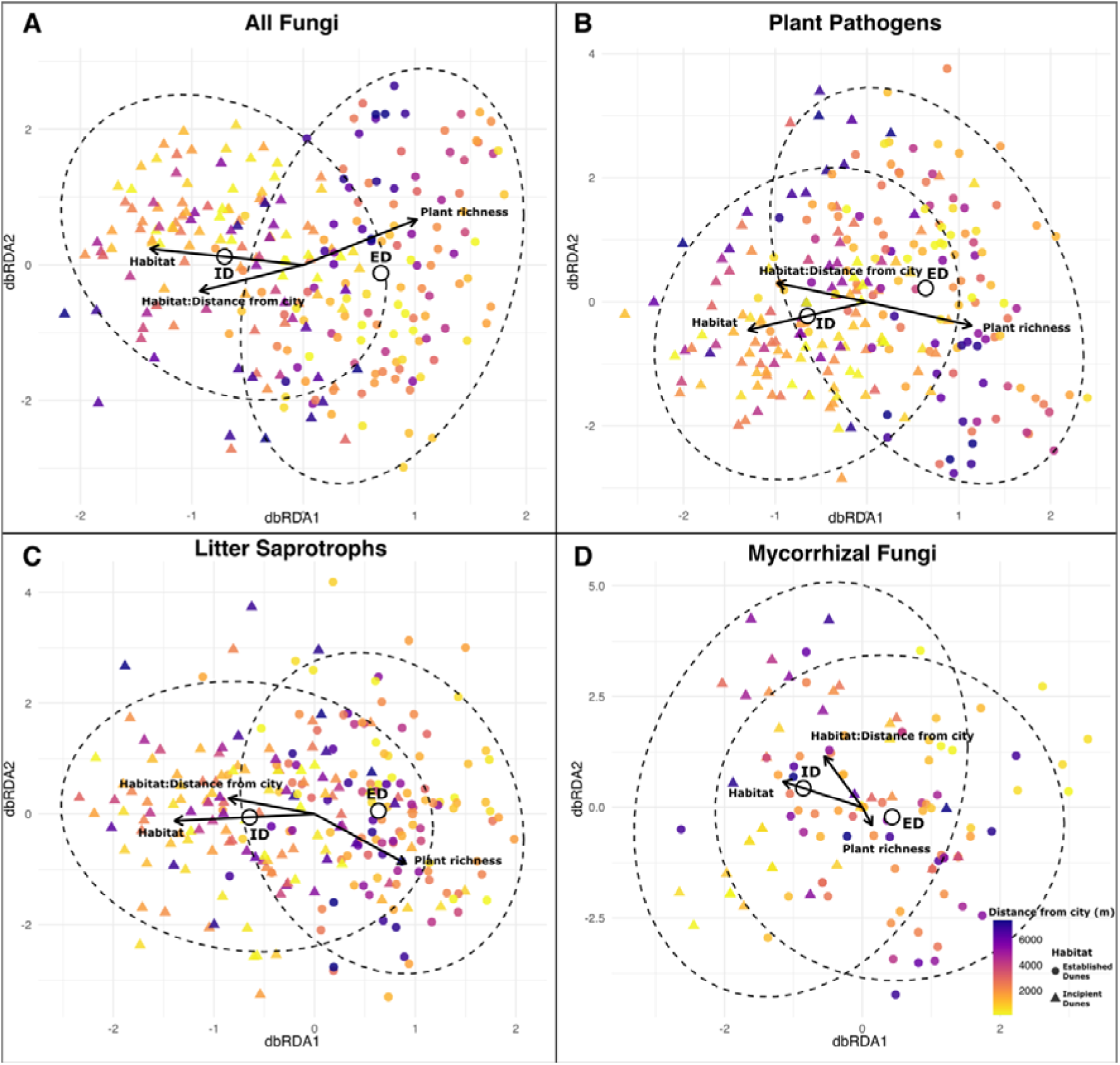
Distance-based redundancy analysis (dbRDA) of fungal communities in coastal dunes along an urbanisation gradient. Bray–Curtis dissimilarities are based on HellingerLtransformed OTU abundances. Points are samples, coloured by distance from the nearest city and shaped by habitat (triangles = incipient dunes, circles = established dunes); ellipses show 95% data ellipses. Arrows are biplot scores for predictors (sediment grain size, plant richness), with length indicating relative importance and direction the gradient of increasing values; grain size is included in the model but its arrow is omitted because it explained no additional variation. Distance from the city is constant within transects and, because transect is controlled for, its effect appears only through the dune type × distance interaction rather than as a separate arrow. Panels: (A) all fungi, (B) plant pathogenic fungi, (C) litter saprotrophic fungi, (D) mycorrhizal fungi.

For plant pathogenic fungi, multi-site β-diversity partitioning (Sørensen) showed that turnover dominated compositional differences (β_SIM = 0.984; 99.4% of total β_SOR = 0.990), with nestedness contributing only 0.6% (β_SNE = 0.006). After controlling for transect, the dbRDA based on Bray–Curtis dissimilarities was significant (F = 8.85, p = 0.001). The constrained component explained 10.5% of the variation in pathogen community composition (adjusted R² = 0.119). Marginal tests indicated significant effects of plant richness (F = 5.49, p = 0.001) and of the dune type × distance-to-city interaction (F = 2.58, p = 0.004), consistent with habitat-specific responses to urbanisation, whereas sediment grain size did not explain additional variation. The first two constrained axes (CAP1 and CAP2) were highly significant (both p = 0.001) and together accounted for 95.3% of the constrained variance (81.8% and 13.5%, respectively), whereas CAP3 was not significant (p = 0.155; Figure 2B).

In litter saprotrophs, multi-site β-diversity (Sørensen) was high (β_SOR = 0.989) and driven almost entirely by turnover (β_SIM = 0.982; 99.2% of total β), with only a small contribution of nestedness (β_SNE = 0.008; 0.8%). After controlling for transect, the dbRDA based on Bray–Curtis dissimilarities was significant (F = 12.05, p = 0.001). The constrained component explained 13.5% of the variation in saprotroph community composition (adjusted R² = 0.158). Marginal tests indicated significant effects of plant richness (F = 3.22, p = 0.003) and of the dune type × distance-to-city interaction (F = 3.10, p = 0.003), consistent with habitat-specific responses along the urbanisation gradient, whereas sediment grain size did not explain additional variation. The first two constrained axes (CAP1 and CAP2) were significant (p = 0.001 and p = 0.012, respectively) and together accounted for 95.7% of the constrained variance (89.0% and 6.7%, respectively); CAP3 was weaker but still significant (p = 0.029; Figure 2C).

For mycorrhizal fungi, β-diversity (Sørensen) was high (β_SOR = 0.987) and dominated by turnover (β_SIM = 0.979; 99.2% of total β), with only a small contribution of nestedness (β_SNE = 0.008; 0.8%). After controlling for transect, the dbRDA based on Bray–Curtis dissimilarities was significant (F = 2.51, p = 0.001). The constrained component explained 5.7% of the variation in mycorrhizal community composition (adjusted R² = 0.056). Marginal tests indicated no clear effect of plant richness (F = 1.44, p = 0.120), and sediment grain size did not explain additional variation, whereas the dune type × distance-to-city interaction was significant (F = 2.04, p = 0.007), pointing to habitat-specific responses to urbanisation. Among the constrained axes, CAP1 was significant (p = 0.001) and accounted for 64.7% of the constrained variance, whereas CAP2 and CAP3 were not significant (p = 0.27 and p = 0.29, respectively) and together contributed a further 20.2% (Figure 2D).

### Local contributions to beta diversity (LCBD)

Across all fungi, LCBD values were low but non-zero (mean ± SD ≈ 0.0040 ± 0.00048; range 0.0030–0.0062) and very similar between dune types. LCBD declined with increasing distance from the city in both habitats, with a stronger decline in incipient dunes than in established dunes (established dunes: edf = 1.71, χ² = 5.90, p = 0.025; incipient dunes: edf = 1.88, χ² = 33.10, p < 0.001). Sediment grain size had no detectable effect on LCBD (p = 0.76), and plant richness showed only a weak, non-significant association (edf = 1.82, χ² = 5.76, p = 0.081). The model explained 43.1% of the deviance (adjusted R² = 0.342), and the random effect for transect indicated substantial among-transect heterogeneity in site uniqueness (Figure 3A).

**Figure 3:**
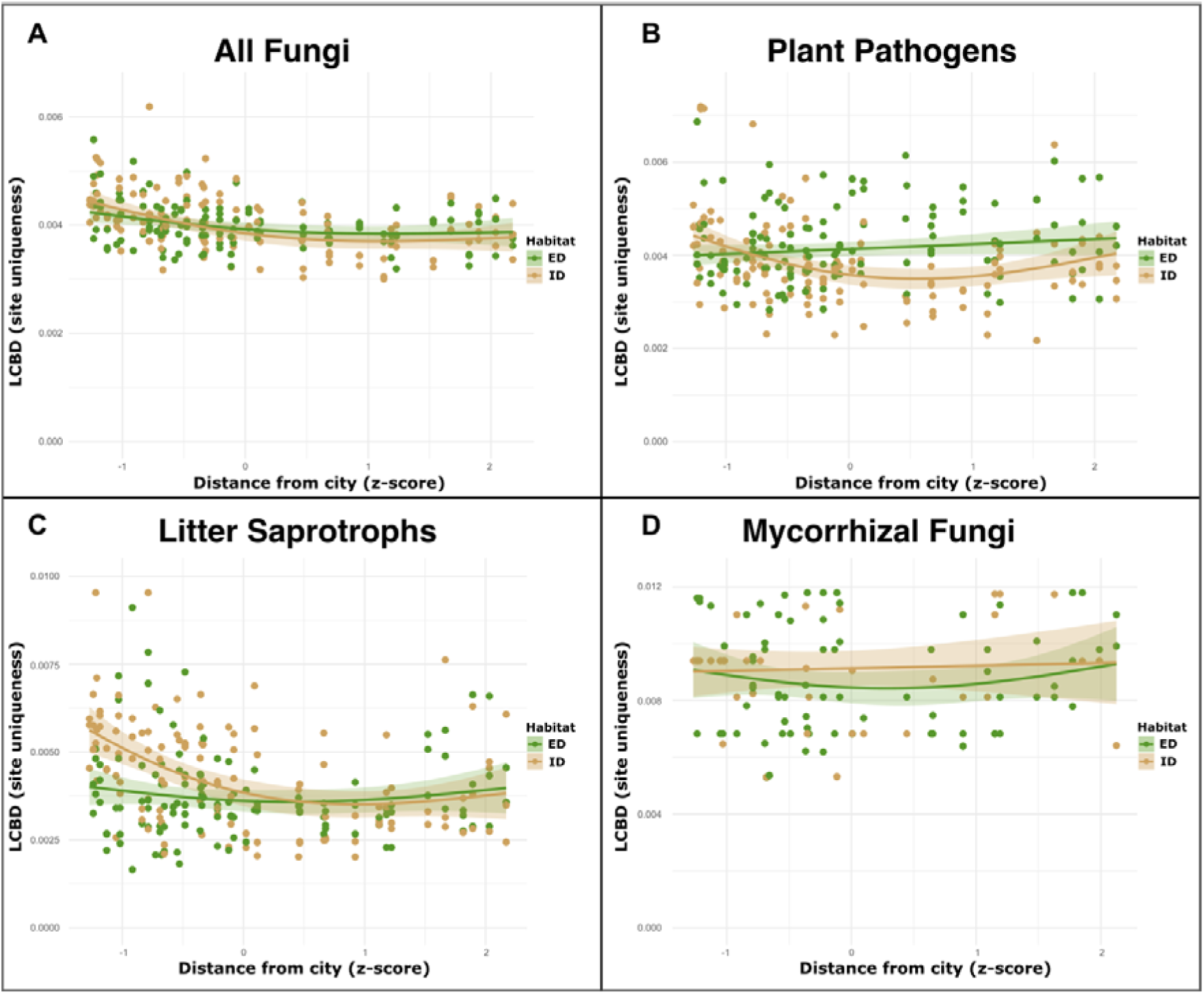
Community uniqueness (LCBD) in coastal dune habitats with increasing distance from cities. Points show site LCBD values computed from Hellinger-transformed OTU data. Lines are GAM (beta regression) fits with 95% confidence ribbons, shown separately for established dunes (ED) and incipient dunes (ID). Distance from the nearest city is z-scored; grain size and plant richness are held at their mean values for predictions. Panels: (A) all fungi, (B) plant pathogenic fungi, (C) litter saprotrophic fungi, (D) mycorrhizal fungi.

For plant pathogenic fungi, LCBD values were generally low but non-zero (mean ± SD ≈ 0.0040 ± 0.00089; range 0.0022–0.0072) and were slightly higher in established dunes than in incipient dunes (0.00423 ± 0.00081 vs. 0.00383 ± 0.00093). LCBD varied with distance from the city only in incipient dunes (edf = 1.92, χ² = 18.16, p < 0.001), showing a non-linear pattern with a minimum at intermediate distances, while no clear trend with distance was detected in established dunes (edf ≈ 1.00, χ² = 2.14, p = 0.14). Both sediment grain size and plant richness contributed significantly to variation in LCBD (grain size: edf ≈ 1.54, χ² = 9.55, p = 0.004; plant richness: edf ≈ 1.00, χ² = 11.64, p < 0.001). The model explained 26.3% of the deviance (adjusted R² = 0.226); the transect random effect was weak and not clearly supported (Figure 3B).

For litter saprotrophs, LCBD values were generally low but variable across sites (mean ± SD ≈ 0.0041 ± 0.0014; range 0.0017–0.0095) and were slightly higher in incipient dunes than in established dunes (0.00435 ± 0.00150 vs. 0.00378 ± 0.00127; main effect of dune type, p = 0.010). LCBD varied with distance from the city only in incipient dunes, with a strong non-linear effect of distance (edf ≈ 1.90, χ² = 30.23, p < 0.001), whereas no clear trend was detected in established dunes (edf ≈ 1.66, χ² = 1.60, p = 0.38). Sediment grain size and plant richness had no detectable effects on LCBD (both p > 0.5). The model explained 34.3% of the deviance (adjusted R² = 0.258), and the transect random effect indicated marked among-transect heterogeneity in site uniqueness (Figure 3C).

For mycorrhizal fungi, site uniqueness (LCBD) was modest but variable across sites (mean ± SD ≈ 0.0089 ± 0.0018; range 0.0053–0.0118) and was similar between established and incipient dunes (0.00881 ± 0.00191 vs. 0.00893 ± 0.00169; main effect of dune type, p = 0.30). LCBD showed no relationship with urbanisation in either habitat (established dunes: edf ≈ 1.61, χ² = 1.56, p = 0.47; incipient dunes: edf ≈ 1.00, χ² = 0.10, p = 0.75). Grain size and plant richness showed no clear effects on LCBD (p ≈ 0.07 and p = 0.18, respectively), and the transect random effect accounted for little additional variation (p = 0.81). Overall model fit was low (deviance explained = 8.8%, adjusted R² = 0.026; Figure 3D).

## Discussion

Urbanisation reshapes dune fungal communities, but not uniformly. Rather than simply reducing richness, it acts as an environmental filter that replaces taxa based on traits, producing non-linear responses and, for most guilds, higher site uniqueness near cities. Across alpha, beta and LCBD analyses, responses are clearly habitat-and guild-specific: effects are generally strongest in incipient (early-successional) dunes, whereas established dunes show weaker or buffered responses. These patterns support our hypotheses that urbanisation impacts are strongest in incipient dunes, that functional guilds respond differently, and that sites near cities harbour more compositionally unique fungal assemblages. Together, they clarify when and where urban pressures most strongly reorganise soil fungal biodiversity in coastal dunes.

### Whole-community response: stronger urban effects in incipient dunes

At the whole-community level, incipient and established dunes differed in composition, but alpha diversity did not differ significantly between habitats once plant richness was accounted for. This is consistent with dune succession, where vegetation develops and plant-fungal coupling strengthens with soil formation and increasing cover (Roy-Bolduc et al., 2016a), and suggests that vegetation context explains much of the habitat contrast in fungal diversity. Even so, the lower vegetation cover and richness, mobile sands, and greater human disturbance (e.g. trampling, vegetation removal near infrastructure) characteristic of incipient dunes likely still contribute to the observed compositional differences.

These contrasting responses reflect fundamental differences in environmental stability and buffering capacity between habitats. Established dunes, with greater geomorphic stability, more complex vegetation, and buffered microclimates, can mitigate disturbance impacts on plant diversity (Anthony Stallins, 2003; Miller et al., 2010), and similar buffering has been suggested for fungi (Brown, 1958; Prenafeta Boldú et al., 2014). In contrast, stronger responses in incipient dunes likely stem from their early-successional state and exposure to multiple, compounding stressors such as low vegetation cover, mobile sand, fluctuating microclimates, and high disturbance. Even stress-tolerant fungi (e.g. salt-or desiccation-resistant taxa) may fail to colonise or persist when urban pressures are added on top of these conditions, indicating limits to physiological tolerance under combined filtering (Rillig et al., 2023). In this context, the more stable vegetation and microclimate of established dunes likely reduce trait–environment mismatches and help maintain fungal community integrity under moderate urban influence.

### Non-linear community responses and the role of multiple stressors

In incipient dunes, fungal diversity showed a non-linear relationship with urban proximity: diversity was lowest at intermediate distances and higher both close to cities and at remote sites, whereas established dunes changed little with distance. This pattern suggests sensitivity to multiple, interacting stressors. Similar non-linear responses to urbanisation have been observed in other ecosystems (Whitehead et al., 2022; Rusterholz and Baur, 2023), where disturbance-tolerant generalists dominate urban-proximate sites and specialists thrive in more natural areas. The intermediate parts of the gradient may act as “mismatch zones”, where trait–environment combinations are suboptimal and neither group performs well. Previous work also shows that specific fungal groups, guilds and habitats are especially vulnerable to disturbance and environmental filtering (Epp Schmidt et al., 2017),(Hall et al., 2009),(Eldridge et al., 2021), which is consistent with the patterns we observe.

The combination of non-linear responses and elevated uniqueness of sites near cities is expected when multiple environmental filters act simultaneously and interact non-additively along the gradient. Several mechanisms are likely involved. Physical disturbance (trampling, sediment movement, vegetation loss) disrupts hyphal networks and disproportionately affects fungi with extensive below-ground structures (Hannula et al., 2017),(Whitehead et al., 2022; Rusterholz and Baur, 2023). Nutrient enrichment from urban inputs can alter competition, favouring fast-growing species and destabilising mutualisms adapted to nutrient-poor dune soils (Hall et al., 2009). Microclimatic change can filter out taxa with narrow tolerances(Khalid et al., 2024), and turnover in host communities, including the presence of non-native plants, alters available niches, potentially facilitating some pathogens while reducing mutualistic interactions (da Silva et al., 2015),(Canini et al., 2019).

### Community turnover and compositional reassembly

Beta□diversity analyses showed that fungal communities are strongly structured by dune habitat, with additional habitat□specific responses along the urbanisation gradient and an important contribution of plant richness for most guilds. Turnover, rather than nestedness, drove compositional shifts, consistent with taxon replacement under trait□based environmental filtering and in line with previous work on fungi (Masumoto et al., 2023),(Hannula et al., 2019). This dominance of turnover indicates that urbanisation and habitat differences do not simply erode richness, but favour different taxon sets, leading to compositional reassembly across both natural (habitat type) and anthropogenic (urban proximity) gradients (Cho et al., 2017). Plant richness explained additional variation in composition for most groups, underlining the role of vegetation in structuring fungal communities. Importantly, habitat × distance-from-city effects remained after accounting for plant richness, indicating that urban filters act in addition to variation in plant richness.

Patterns of community uniqueness (LCBD) reinforce these findings. Across all fungi, LCBD declined with distance from the city, with non-linear responses in both habitats and a stronger decline in incipient dunes. Plant richness showed only a weak, non-significant association with LCBD, and grain size had little to no effect. At the guild level, LCBD varied with distance only in incipient dunes for plant pathogens and litter saprotrophs, while mycorrhizal LCBD showed no relationship with distance. Predicted LCBD values tended to be higher near cities, especially in incipient dunes, suggesting that these sites host communities that are compositionally atypical relative to the regional pool. This higher uniqueness near cities likely arises because established dunes buffer disturbance through greater vegetation cover and soil development, whereas incipient dunes remain more open to colonisation by urban-tolerant taxa or stochastic colonisation events.

### Guild-specific responses

Rather than simply reducing richness, urbanisation acts as an environmental filter that replaces taxa based on traits, producing clear guild-specific patterns. These divergent responses confirm our expectation that different fungal groups respond differently to urbanisation, and suggest that outcomes depend on particular trait combinations that either confer advantages or create vulnerabilities under urban pressures (Aguilar-Trigueros et al., 2025).

Litter saprotrophs showed strong filtering by urbanisation, especially in incipient dunes. Near cities, reduced or altered input of organic material (e.g. through wrack removal, lower vegetation due to trampling or buildings) and modified microclimates likely favour disturbance-tolerant, fast-colonising generalists, leading to assemblages that are compositionally unique but poorer in species (Gómez-Hernández et al., 2021),(Semchenko et al., 2022). Consistent with this, Shannon diversity increased with distance from cities in both habitats, whereas community uniqueness was higher close to cities. Further from cities, more stable vegetation and organic litter inputs likely support habitat-associated specialists, matching the dominance of turnover rather than directional loss. If beach management continues to reduce organic inputs, decomposer networks may simplify and carbon cycling may be altered. Functional redundancy among decomposers may buffer some functions (Banerjee et al., 2016),(Veen et al., 2021), but management that retains wrack, limits grooming and trampling, and maintains vegetation should be a priority to sustain more typical saprotroph communities and associated processes.

Plant pathogenic fungi patterns suggest a host-and disturbance-mediated filtering that is strongest in incipient dunes. In these habitats, both alpha diversity and LCBD show minima at intermediate distances from cities, hinting at the mixing of two pathogen assemblages: one associated with urban environments (e.g. spillover from ornamental plants, hosts thriving under disturbed conditions, altered microclimates, and higher fertiliser or pollutant inputs), and one associated with more stable, native vegetation under lower urban influence (Semchenko et al., 2022),(Morriën et al., 2017). Intermediate sites may represent a trait–environment mismatch zone where host communities are mixed or sparse, reducing available niches for pathogens. In contrast, established dunes, with greater vegetation cover, appear to buffer pathogen communities even close to cities. That plant richness explained additional compositional variation in the dbRDA and also influenced LCBD is consistent with pathogens tracking host availability: sites with richer host communities can support pathogen assemblages that are both compositionally distinct and more unique, suggesting roles for host identity, density, and total richness (Rottstock et al., 2014; Rutten et al., 2021). The dominance of turnover over nestedness indicates taxon replacement rather than directional loss, consistent with urban filters favouring disturbance-tolerant taxa while remote dunes favour specialists linked to native hosts. If disturbance and ornamental plants introduce novel host–pathogen pools, managers should limit non-native plant introductions, reduce nutrient enrichment and physical disturbance near foredunes, and monitor for pathogen spillover into native vegetation. Future work should link pathogens to host identity and density, quantify disease incidence along the gradient, and test experimentally whether urban-proximate incipient dunes are hotspots for pathogen establishment and transmission.

Mycorrhizal fungi alpha diversity in incipient dunes increased from close to cities towards intermediate distances, consistent with recovery or higher availability of compatible hosts, and with known traits of many mycorrhizal taxa that have low dispersal capacity and high disturbance sensitivity (Chaudhary et al., 2022),(Epp Schmidt et al., 2017). However, the absence of strong peaks in community uniqueness suggests more diffuse shifts that are not tightly linked to distance from a city. This pattern is compatible with strong host dependence, where mycorrhizal fungi primarily track host availability wherever suitable hosts occur, rather than distance per se. The weak effects of grain size and the non-significant effect of plant richness support the idea that host identity, rather than total plant richness, is the main constraint, as reported in other systems (Martínez-García et al., 2015; Matsuoka et al., 2020). In established dunes, stable vegetation and soils likely buffer mycorrhizal assemblages against urban influences, although slight richness changes may still reflect variation in host availability near cities. Maintaining or restoring native host plants and preserving source populations of both hosts and fungi should facilitate mycorrhizal colonisation and diversity in dunes. Future work could quantify colonisation speed and host specificity, and test whether reduced urban disturbance accelerates the diffuse turnover observed here into more fully restored, host-matched mycorrhizal communities.

### Future directions

Although we did not measure ecosystem processes directly, our results suggest clear hypotheses. Reduced saprotroph richness and strong compositional turnover near cities may alter decomposition and nutrient cycling unless functional redundancy buffers these functions. Reassembly of mycorrhizal communities could influence host establishment and dune stabilisation, while turnover in plant pathogens may shift disease pressure depending on host identity and availability.

A key priority for future work is to link species-level traits to community change, and moving from describing patterns to understanding mechanisms. Integrating traits such as dispersal capacity, symbiotic dependence, stress tolerance and competitive ability into trait-environment analyses (e.g. fourth-corner tests or trait-informed joint species distribution models) would allow explicit tests of whether taxon replacement reflects trait divergence, and improve predictions along the urban gradient (Camenzind et al., 2024),(Chaudhary et al., 2022). These trait data should be paired with information on host plant composition and traits, as well as soil properties, to assess how fungal reassembly tracks vegetation change and nutrient regimes. Because our study provides only a spatial snapshot, denser sampling across broader spatial and temporal scales is needed to separate short-term from persistent responses, track turnover and community uniqueness through time, and detect legacy effects of disturbance. Manipulative experiments that vary nutrient inputs, physical disturbance, vegetation structure and microclimate could identify causal filters and their interactions, as shown in other habitats (Cho et al., 2017; Fanin et al., 2022; Zhan et al., 2023). Finally, coupling community profiling with process measurements (e.g. decomposition rates, nutrient mineralisation) and functional omics (metatranscriptomics/metaproteomics) would test whether compositional shifts translate into changes in ecosystem functioning (Schneider et al., 2012; Chen et al., 2021; Auer et al., 2024), underscoring the value of a trait-based perspective for interpreting community responses to urbanisation.

## Conclusion

Urbanisation restructures dune fungal communities in habitat-and guild-specific ways, with incipient dunes particularly vulnerable to the combined effects of urban disturbance and natural stressors. Rather than simply reducing richness, urbanisation acts as a trait-based filter that replaces taxa and often increases community uniqueness near cities, especially for saprotrophic and plant pathogenic fungi. Established dunes show weaker, likely more buffered responses, underscoring the importance of habitat context in the coastal dune ecotone. Together, these results clarify when and where urban pressures most strongly reorganise soil fungal biodiversity in coastal dunes and highlight the need to prioritise protection and restoration of early-successional dunes by limiting disturbance and maintaining host plant availability.

## Data Availability Statement

All raw data is available in the NCBI Sequence Read Archive (SRA), under BioProject number PRJNXXXX

## Acknowledgements

We would like to thank the Naturalis lab team for their support, and Rijkswaterstaat, Staatsbosbeheer, Dunea, and Waternet for facilitating access to the study area. This study is part of the MeioMon grant to JNM, funded by the Stemmler Foundation.

